# Advancing CAR T-Cell Therapy: Simultaneously Attack Tumor and Immunosuppressive Cells in the Tumor Microenvironment

**DOI:** 10.1101/2024.03.25.586653

**Authors:** Yan Luo, Emiliano Sanchez Garavito, Mieu M. Brooks, Martha E. Gadd, Yaqing Qie, Maria J. Ulloa Navas, Tanya Hundal, Shuhua Li, Andrea Otamendi Lopez, Vanessa K. Jones, Yanyan Lou, Tushar Patel, Roxana Dronca, Mohamed A. Kharfan-Dabaja, Haidong Dong, Alfredo Quinones-Hinojosa, Hong Qin

**Author notes:** Correspondence: Hong Qin, MD, PhD Associate Professor of Medicine, Division of Hematology/Oncology Director, Regenerative Immunotherapy & CAR-T Translational Research Program Mayo Clinic Florida 4500 San Pablo Rd S, Jacksonville, FL 32224.

## Abstract

Chimeric antigen receptor (CAR) T-cell therapy has encountered limited success in solid tumors. The lack of dependable antigens and the immunosuppressive tumor microenvironment (TME) are major challenges. Within the TME, tumor cells along with immunosuppressive cells employ an immune-evasion mechanism that upregulates programmed death ligand 1 (PD-L1) to deactivate effector T cells; this makes PD-L1 a reliable, universal target for solid tumors. We developed a novel PD-L1 CAR (MC9999) using our humanized anti-PD-L1 monoclonal antibody, designed to simultaneously target tumor and immunosuppressive cells. The antigen-specific antitumor effects of MC9999 CAR T-cells were observed consistently across four solid tumor models: breast cancer, lung cancer, melanoma, and glioblastoma multiforme (GBM). Notably, intravenous administration of MC9999 CAR T-cells eradicated intracranially established LN229 GBM tumors, suggesting penetration of the blood-brain barrier. The proof-of-concept data demonstrate the cytolytic effect of MC9999 CAR T-cells against immunosuppressive cells, including microglia HMC3 cells and M2 macrophages. Furthermore, MC9999 CAR T-cells elicited cytotoxicity against primary tumor-associated macrophages within GBM tumors. The concept of targeting both tumor and immunosuppressive cells with MC9999 was further validated using CAR T-cells derived from cancer patients. These findings establish MC9999 as a foundation for the development of effective CAR T-cell therapies against solid tumors.

## Introduction

Immunotherapies are cancer treatments that harness the immune systems of patients to fight their diseases. Two well-known examples are immune checkpoint inhibitors (ICIs) and chimeric antigen receptor (CAR) T-cells. ICIs target inhibitory or stimulatory pathways that modulate immune cell activity, with many ICIs specifically mobilizing cytotoxic T cells.^1,2^ CAR T-cells are a personalized therapy in which patient T cells are engineered to express a membrane spanning receptor with an extracellular domain that recognizes a cancer antigen and an intracellular domain that activates the T cell leading to an antigen-specific cellular therapy.^3,4^

CAR T-cells have been revolutionary in the treatment of relapsed/refractory B cell malignancies due to the antigen specificity to the target tissue, namely CD19, as well as the medical management of B cell aplasia.^5^ However, identification of such a target antigen is a challenge for solid tumors, particularly when considering that the presence of antigen heterogeneity complicates finding a universal antigen for CAR T-cell therapy. In addition, solid tumor cells have developed complex interactions with surrounding stromal cells while recruiting and enlisting diverse immune cells, to create an immunosuppressive tumor microenvironment (TME), allowing the tumor cells to escape antitumor immunity.^6,7^

A potential target that can address the hurdles of antigen heterogeneity and immunosuppression is programmed death-ligand 1 (PD-L1). The PD-1/PD-L1 axis functions natively as a regulator of immune tolerance and as a governor for T cell activation. Cancer has manipulated this inhibitory cascade to evade T cell attack. Upregulation of PD-L1 expression on various solid tumors has been well-documented,^2,8^ leading to FDA approval of ICIs targeting PD-L1 for a diverse collection of solid tumors. This same T cell inhibitory mechanism can also be employed by those immunosuppressive cells within the TME, which is supported by the findings of elevated level of PD-L1 on tumor-associated macrophages (TAMs), myeloid-derived suppressor cells (MDSCs) and regulatory T cells (Tregs).^9–11^ These findings highlight PD-L1 as a promising target for development of a CAR T-cell therapy against both tumor cells and the PD-L1-expressing immunosuppressive cells.

In this study, we developed a PD-L1 CAR T-cell therapy, designated MC9999, using a humanized anti-human PD-L1 monoclonal antibody. We demonstrated the anti-tumor effects of MC9999 CAR T-cells against various solid tumor models, including patient-derived primary tumor cell lines. Intravenously dosed MC9999 CAR T-cells exhibited robust in vivo anti-tumour efficacy and long-term survival in xenograft mouse models of intramammary triple-negative human breast cancer and intracranial glioblastoma multiforme (GBM). We also showed antigen-specific cytotoxicity of MC9999 CAR T-cells against three immunosuppressive cell models, including primary TAMs. To underscore the translational application of MC9999 CAR T-cells, our work culminated in the successful generation of GBM patient-derived MC9999 CAR T-cells that showed cytotoxicity against primary GBM tumor cells and tumor-associated microglial cells. This composite of work establishes PD-L1-targeting MC9999 CAR T-cells as a promising immunotherapy with therapeutic application in solid tumors.

## Results

### Antigen-specific cytotoxicity of anti-PD-L1 CAR T-cells

Using the variable regions of the heavy and light chains of a humanized PD-L1 monoclonal antibody generated by Dr. Dong,^12^ we created the novel MC9999 CAR, which is a second-generation CAR construct that contains the 4-1BB costimulatory domain and the CD3ζ signaling domain. Additionally, we included a truncated epidermal growth factor receptor (tEGFR; Refer to Figure S1) as a safety switch. Our validated manufacturing procedures ensured the quality of CAR T-cell production, resulting in reproducible batches of MC9999 CAR T-cells that met specified quality control measurements (Refer to Table S1).^13^

Antigen-specific cytotoxicity of MC9999 CAR T-cells was confirmed against a PD-L1 overexpressing human breast cancer cell line (MDA-MB-231 PD-L1 OE) with a PD-L1 knock out variant (MDA-MB-231 PD-L1 KO) as a negative control. Non-transduced T cells (Non-CAR T-cells) from the same donor were used as an alloreactivity control. Both CD8 and CD4 MC9999 CAR T-cells exhibited cytotoxicity in response to MDA-MB-231 PD-L1 OE, as evidenced by T-cell degranulation with subsequent surface expression of CD107a. The absence of cytotoxic activity of the same CAR T-cell populations to MDA-MB-231 PD-L1 KO affirmed the antigen-specific functionality (Figure 1A and Figure 1B, respectively). Consistently, a significant release of granzyme B was observed when MC9999 CAR T-cells were incubated with MDA-MB-231 PD-L1 OE but not with MDA-MB-231 PD-L1 KO (Figure 1C). Antigen-specific cytolysis of MC9999 CAR T-cells was further evaluated from the perspective of the target cells in an impedance-based killing assay. Cytolysis, as determined with the decrease in the cellular electrical impedance of cultured target cells, was observed only in MDA-MB-231 PD-L1 OE cells (Figure 1D); all other conditions showed that target cells remained intact.

**Figure 1.**
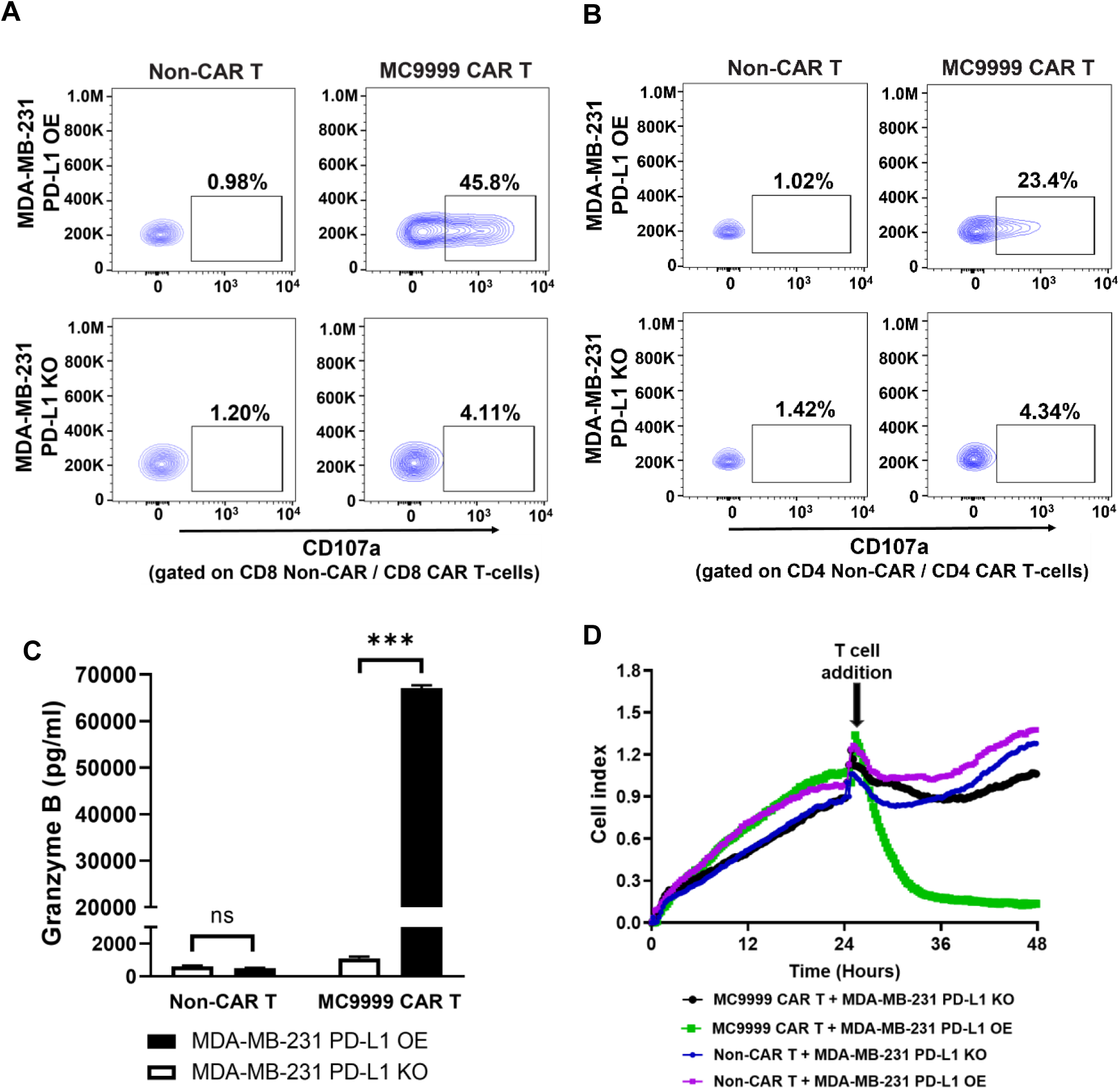
MC9999 CAR T-cells showed antigen-specific cytotoxicity against PD-L1 expressing breast cancer cells. CAR T-cell cytotoxicity was evaluated by CD107a surface expression in a degranulation assay. MC9999 CAR T-cells were incubated with MDA-MB-231 PD-L1 OE or MDA-MB-231 PD-L1 KO cells at an E:T ratio of 2:1. Analysis was gated on CD4+CAR T-cells (A) or CD8+CAR T-cells (B); Non-CAR T from the same donor served as negative controls. (C) MC9999 CAR T-cells were co-cultured with either MDA-MB-231 PD-L1 OE or MDA-MB-231 PD-L1 KO cells. After 72 hours, the supernatants were collected and examined granzyme B by ELISA. The data were plotted as the mean ± SEM of quadruplicate sampling and are representative of three independent experiments (***, p < 0.001; ns, no significance). (D) Target cells, either MDA-MB-231 PD-L1 OE or MDA-MB-231 PD-L1 KO cells, were seeded on the E-plates for 24 hours. After 24 hours, MC9999 CAR T-cells or Non-CAR T-cells were added to the target cells at a E/T ratio of 40:1. Cell index (CI) traces were collected in triplicate every 15 minutes during the co-culture, and changes in impedance were normalized to the 24-hour time point. The results represent three independent experiments.

We next evaluated the in vivo anti-tumor effects of MC9999 CAR T-cells in female NSG mice that received an intramammary challenge of either MDA-MB-231 PD-L1 OE or MDA-MB-231 PD-L1 KO tumor cells, both of which were modified to express luciferase. Mice bearing established tumors were assigned to one of three treatment groups (n=5 per group): MC9999 CAR T-cells, Non-CAR T-cells, or PBS. Using bioluminescence imaging to track tumor progression, we observed a substantial reduction in established MDA-MB-231 PD-L1 OE tumors following treatment with MC9999 CAR T-cells. However, this treatment showed no effect on the growth of MDA-MB-231 PD-L1 KO tumors (Figure 2A), confirming the antigen-specific in vivo antitumor effects. Three Kaplan-Meier plots were generated to compare treatment-associated survival rates between the antigen-positive (MDA-MB-231 PD-L1 OE) and the antigen-deficient (MDA-MB-231 PD-L1 KO) tumor challenge groups [treatment groups: MC9999 CAR T-cells (Figure 2B), Non-CAR T-cells (Figure 2C), and PBS (Figure 2D)]. Consistent with the bioluminescence imaging findings, MC9999 CAR T-cell treatment resulted in prolonged survival exceeding 120 days in mice challenged with MDA-MB-231 PD-L1 OE tumor cells, while showing no such antitumor effect in those with MDA-MB-231 PD-L1 KO tumors. Neither Non-CAR T-cell treatment nor PBS demonstrated any antitumor effects, resulting in rapid tumor progression and mortality in all mice within 50 days post-tumor challenge.

**Figure 2.**
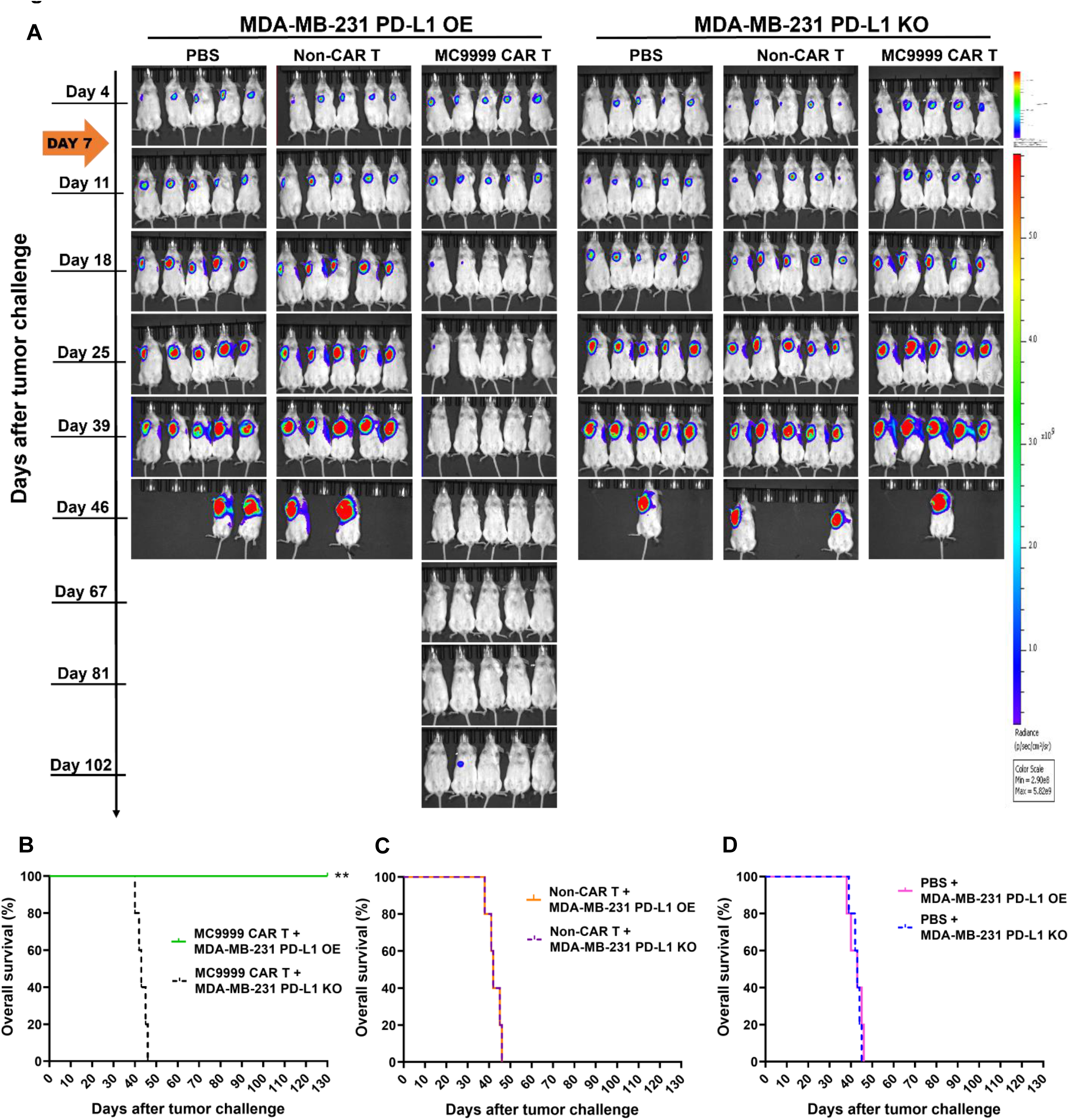
MC9999 CAR T-cells elicited antitumor effects in an intramammary breast cancer model. (A) Female NSG mice were given an intramammary injection of luciferase-labeled MDA-MB-231 tumor cells with either PD-L1 overexpressing (PD-L1 OE) or PD-L1 knockout (PD-L1 KO) at a dose of 1.0×10^6^ cells per mouse. Seven days after tumor challenge, mice were randomly divided into three groups. Each group received a single intravenous (IV) infusion of one of the following: PBS, Non-CAR T-cells (5×10^6^ total T cells/per mouse), or MC9999 CAR T-cells (2×10^6^ CAR T-cells out of 5×10^6^ total T cells from the same donor/per mouse). Weekly IVIS^©^ imaging was performed to monitor tumor progression. The representative images demonstrated the changes in tumor burdens over time. The results present two independent experiments using different donor T cells for generating CAR T-cells. Three separate Kaplan-Meier survival plots: MC9999 CAR T-cells (B), Non-CAR T-cells (C), or PBS (D) were generated to evaluate the treatment-associated overall survival. Log-rank analysis revealed that MC9999 CAR T-cell treatment significantly extended the overall survival in mice challenged with MDA-MB-231 PD-L1 OE comparing to those bearing MDA-MB-231 PD-L1 KO tumors (** p = 0.01). No significant differences in survival were observed in the PBS or Non-CAR T-cells treatment group between these two tumor models.

### MC9999 CAR T-cells showed antigen-specific cytotoxicity against various solid tumors

Targeting PD-L1 is supported by reports of PD-L1 expression in a variety of solid tumors such as lung cancer, melanoma, and GBM. As such, we evaluated the cytotoxicity of MC9999 CAR T-cells across these three types of solid tumors. Calu-1, a non-small-cell lung cancer (NSCLC) cell line, stably expresses PD-L1. We generated a PD-L1-deficient variant (Calu-1 PD-L1 KO) (Refer to Figure S2) as a control. In a CD107a T-cell degranulation assay, we observed antigen-specific cytotoxicity of MC9999 CAR T-cells against PD-L1-expressing Calu-1 cells (Figure 3A). The release of granzyme B by the CAR T-cells was exclusively detected in response to Calu-1, but not Calu-1 PD-L1 KO cells, further confirmed the antigen-specific cytotoxicity (Figure 3B).

**Figure 3.**
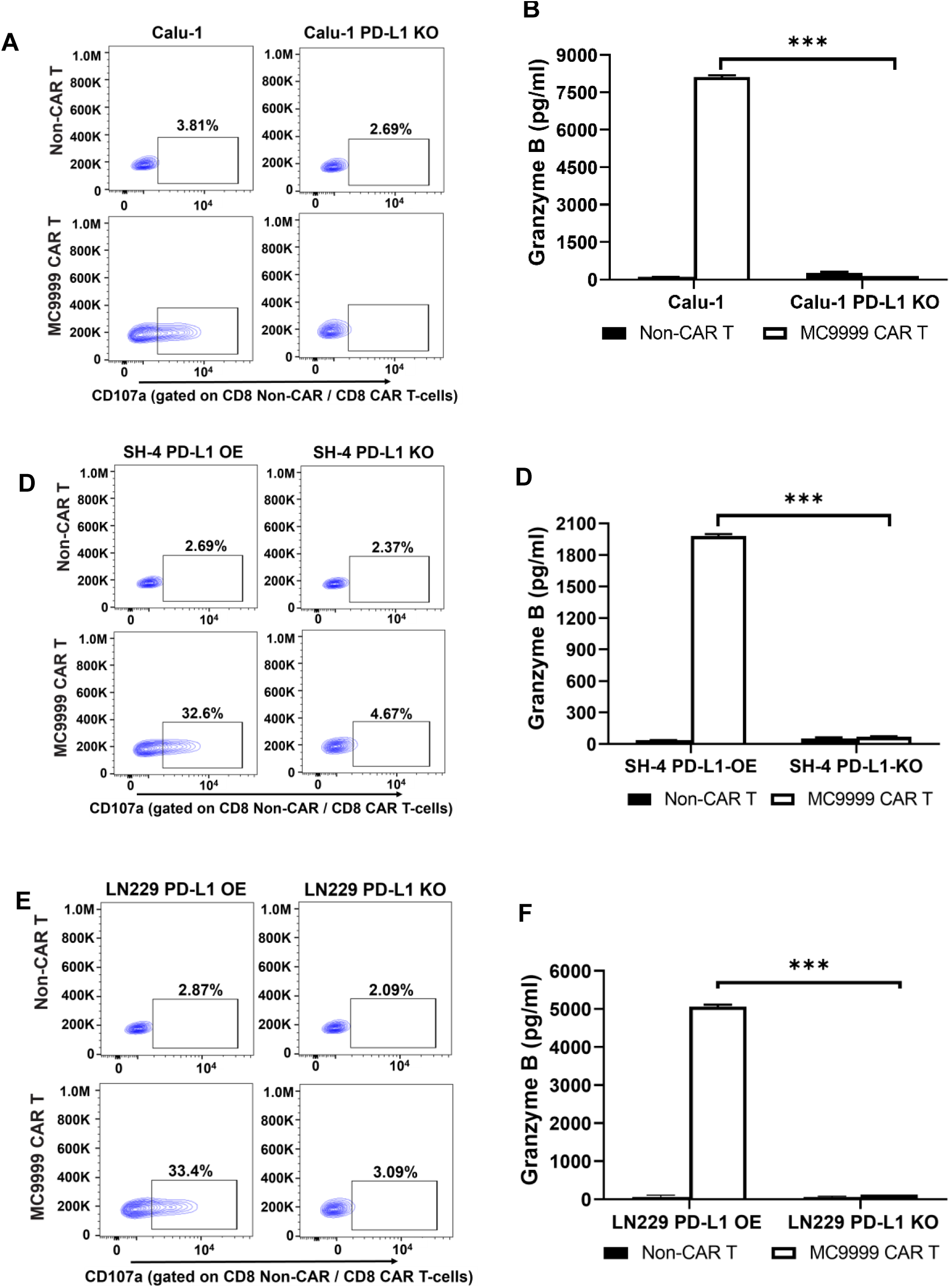
MC9999 CAR T-cells exhibited antigen-specific cytotoxicity against various PD-L1 expressing solid tumors. In a CD107a degranulation assay, antigen-specific cytotoxicity of MC9999 CAR T-cells was assessed against three solid tumor cell lines: Calu-1 lung cancer (A), SH-4 melanoma (C), and LN229 GBM (E). The corresponding PD-L1-deficient tumor cell variants were included as negative controls. The release of granzyme B was readily detected when MC9999 CAR T-cells were exposed to PD-L1-expressing target cells including Calu-1(B), SH-4 (D), and LN229 (F). The data, plotted as mean ± SEM of triplicate sampling, are representative of three independent experiments and analyzed using the multiple T test (*** p < 0.001).

SH-4, a representative metastatic melanoma model, was engineered to generate SH-4 PD-L1 OE (PD-L1-overexpressing) and SH-4 PD-L1 KO (PD-L1-deficient) cell lines (Refer to Figure S2), for usage as target cells to investigate therapeutic effectiveness of MC9999 CAR T-cells against melanoma. Antigen-specific T-cell degranulation was evident when MC9999 CAR T-cells were incubated with SH-4 PD-L1 OE, but not with SH-4 PD-L1 KO cells (Figure 3C). As further confirmation, MC9999 CAR T-cells exhibited a significant release of granzyme B upon interaction with SH-4 PD-L1 OE cells (Figure 3D).

The last tumor model we examined was LN229, a GBM cell line, which was also modified to create antigen-positive LN229 PD-L1 OE and antigen-negative LN229 PD-L1 KO variants. (Refer to Figure S2). Our in vitro assays consistently demonstrated specific cytotoxicity of MC9999 CAR T-cells against LN229 PD-L1 OE cells, evidenced by T-cell degranulation (Figure 3E) and granzyme B release (Figure 3F), whereas the T-cell cytotoxicity was absent against LN229 PD-L1 KO cells. Collectively, these findings provide conclusive evidence for the antigen-specific cytotoxicity of MC9999 CAR T-cells against PD-L1-positive solid tumors.

### MC9999 CAR T-cells exhibited cytotoxicity against patient-derived GBM tumor cells

Traditional cell lines have played crucial roles in laying the foundation for proof-of-principle CAR T-cell development; however, these cell lines are devoid of the characteristics associated with their original microenvironment. Patient-derived tumor cell lines retain the tumor characteristics of the original patient and, likely the clinical response to treatment, making these primary tumor cell lines critical in translational medicine. We have established two GBM patient-derived tumor cell lines (QNS120 and QNS712) from surgically resected GBM tumors (showed in MRI images in Figure 4A), both of which are positive for PD-L1 expression (Figure 4B). MC9999 CAR T-cells exhibited cytotoxicity against QNS120 and QNS712 tumor cells, as observed through T-cell degranulation (Figure 4C) and granzyme B (Figure 4D). To ensure the elicited CAR T-cell cytotoxicity was antigen-specific, we included PD-L1-positive (MDA-MB-231 PD-L1 OE) and PD-L1-negative (MDA-MB-231 PD-L1 KO) controls. The utilization of these patient-derived GBM cell lines confirmed the therapeutic potential of MC9999 CAR T-cells against primary tumor cells.

**Figure 4.**
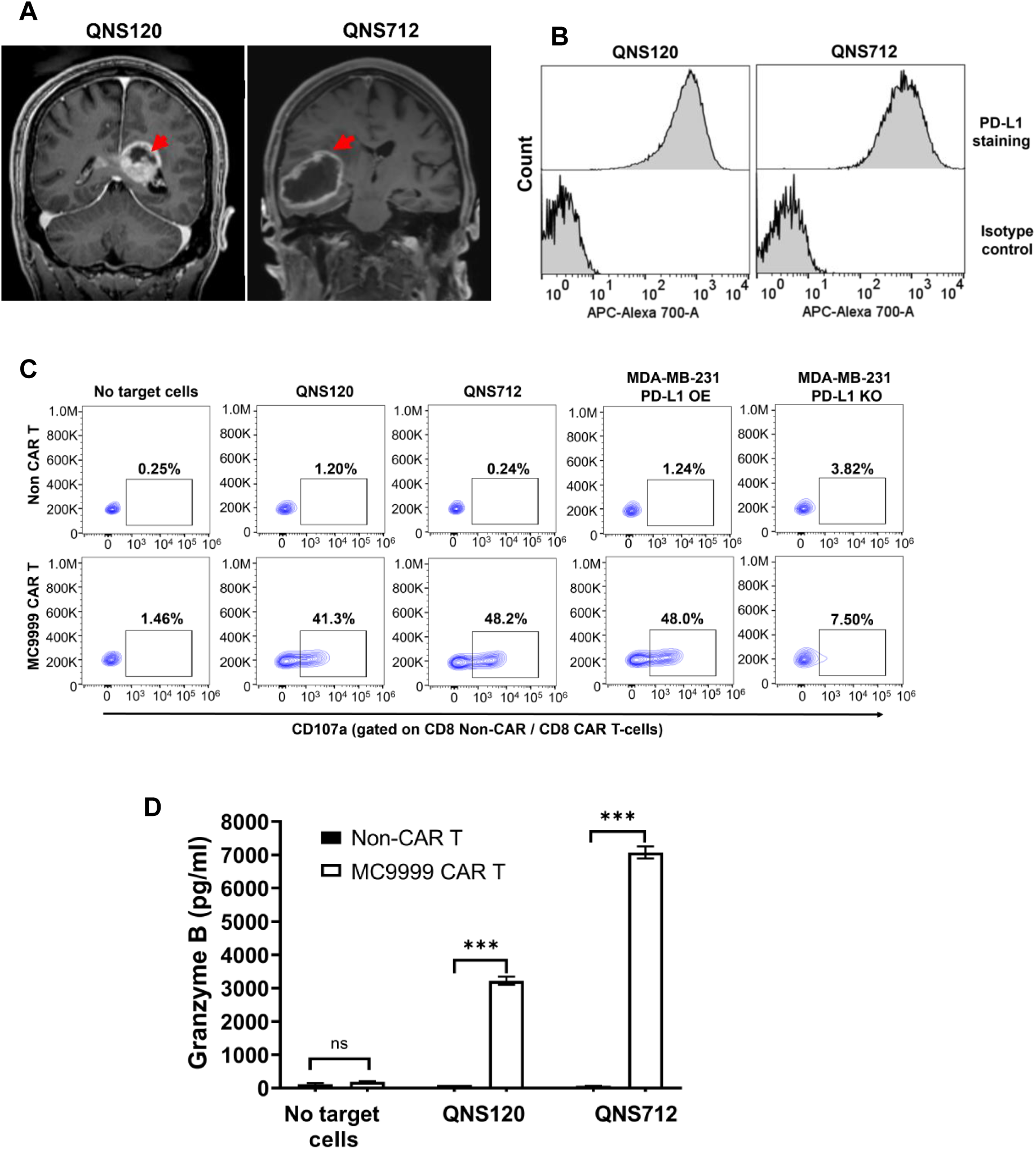
Patient-derived primary GBM cells were targeted by MC9999 CAR T-cells. (A) Two patients (QNS120 and QNS712) were diagnosed with glioblastoma (Grade 4) as confirmed by magnetic resonance imaging (MRI). The red arrow highlights the tumor tissues. (B) The resected tumors were obtained from the patients to generate two patient-derived primary GBM cell lines. The PD-L1 expression on QNS120 and QNS712 tumor cells was characterized using immunostaining and analyzed with flow cytometry. (C) MC9999 CAR T-cells were functionally activated by QNS120 and QNS712 tumor cells, as indicated by cell surface staining of CD107a in a degranulation assay. MDA-MB-231 PD-L1 OE and MDA-MB-231 PD-L1 KO cells were used as antigen-positive and antigen-negative controls, respectively. (D) The release of granzyme B was significantly elevated when MC9999 CAR T-cells targeted QNS120 and QNS712 tumor cells. The data, plotted as mean ± SEM of triplicate sampling, are representative of three independent experiments and analyzed using the multiple T test (*** p < 0.001).

### MC9999 CAR T-cell treatment eradicated intracranially engrafted GBM tumors

We next evaluated the in vivo anti-tumor effects of MC9999 CAR T-cells using the LN229 GBM tumor model (Figure 5A). Mice were challenged with an intracranial injection of luciferase-expressing LN229 PD-L1 OE cells. Treatment with either MC9999 CAR T-cells, Non-CAR T-cells, or PBS was administered intravenously (IV) on Day 7 and Day 14 (orange arrows) following tumor challenge. Bioluminescence images tracked tumor development and revealed significant tumor reduction in mice treated with the MC9999 CAR T-cells (Figure 5A), leading to a substantially extended overall survival. The CAR T-cell therapy enabled these treated mice to achieve tumor-free survival until experiment conclusion on day 150, whereas all mice in PBS and Non-CAR T-cell control groups succumbed to the tumors within 70 days after tumor challenge (Figure 5B). Our findings that IV dosed MC9999 CAR T-cells eradicated intracranially established tumors underscores the capability of these therapeutic T cells to breach and cross the blood-brain barrier.

**Figure 5.**
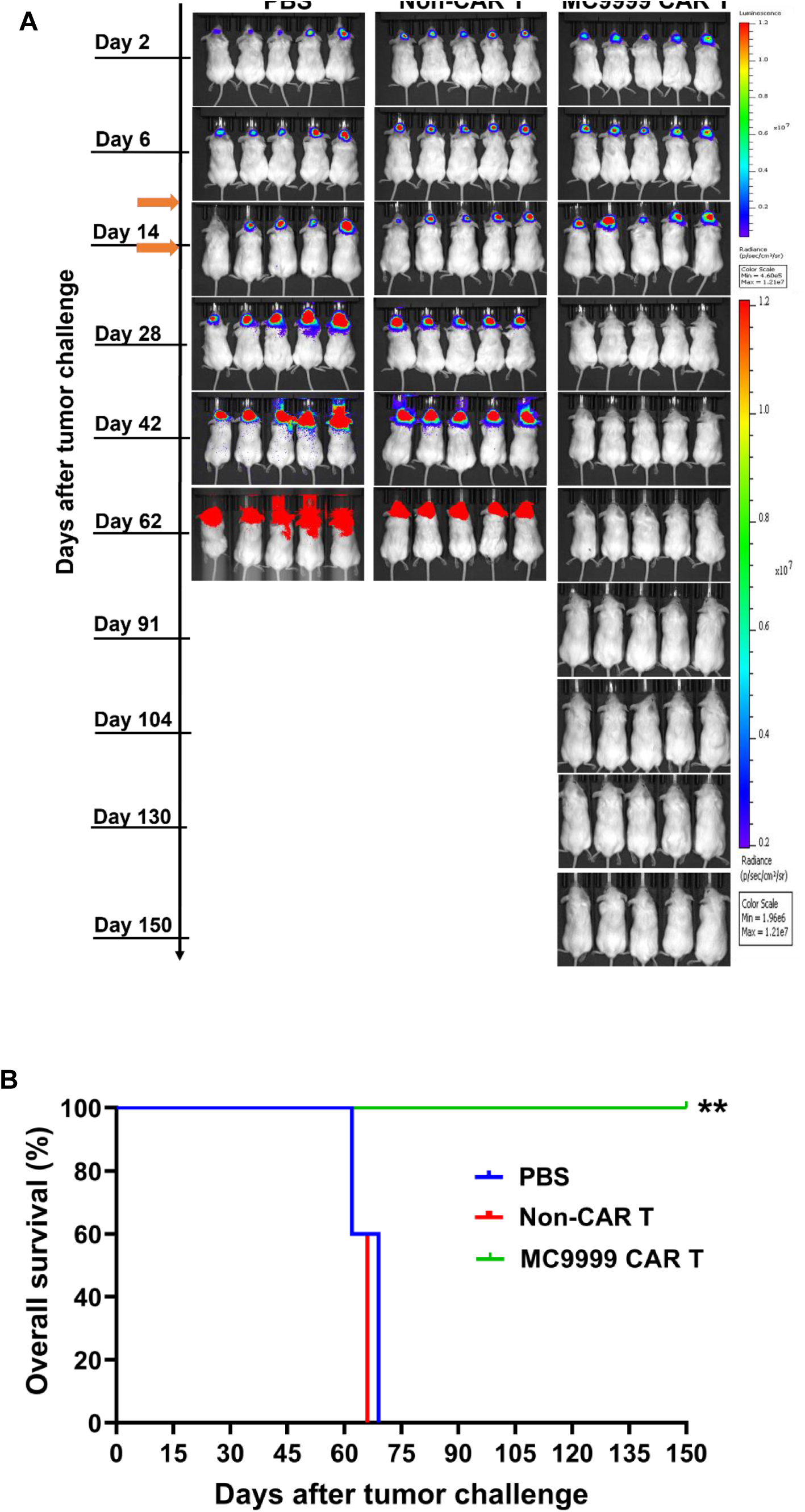
Treatment with MC9999 CAR T-cells eradicated intracranially established LN229 GBM tumors. (A) NSG mice were intracranially challenged with luciferase-labeled, PD-L1-overexpressing LN229 GBM tumor cells at a dose of 0.5×10^6^ cells per mouse. Seven days after the tumor challenge, the mice were randomized into three groups and then received an intravenous (IV) infusion of one of the following: PBS, Non-CAR T-cells (5×10^6^ total T cells/mouse), or MC9999 CAR T-cells (2×10^6^ CAR T-cells out of 5×10^6^ total T cells/mouse) generated from the same donor. A second treatment dose was administrated intravenously on day 14. The mice were imaged weekly to track tumor progression for 150 days. The representative IVIS^©^ images illustrated the changes in tumor burdens over time. (B) A Kaplan-Meier plot was generated to compare the overall survival among the treatment groups. Log-rank analysis revealed significant differences between the MC9999 CAR T-cell treatment group and both control groups receiving either PBS or Non-CAR T-cells (** p < 0.01).

### Immunosuppressive cells as a target for MC9999 CAR T-cells

A major challenge limiting the therapeutic efficacy of CAR T-cell therapies in solid tumor is the TME, that is comprised of various immunosuppressive cells that highjack the PD-1/PD-L1 cascade to inhibit T cell function, allowing tumor cells to evade antitumor immunity.^2,14^ The elevated expression of PD-L1 in immunosuppressive cells makes them a potential target of MC9999 CAR T-cells. We used three immunosuppressive cell models, including HMC3 microglia, monocyte-derived M2 macrophages (MDM-M2), and primary TAMs from GBM patients to test our hypothesis.

Microglia residing within the TME represent a subset of TAMs in GBM.^6^ We first confirmed the expression of PD-L1 in the HMC3 microglial cell line (Figure 6A). Cytotoxicity of MC9999 CAR T-cells against HMC3 cells was evident via CAR T-cell degranulation (Figure 6B) and granzyme B release (Figure 6C). Further evidence was the direct killing of the HMC3 cells upon exposure to MC9999 CAR T-cells that was measured with the disruption of the HMC3 monolayer in an impedance assay (Figure 6D).

**Figure 6.**
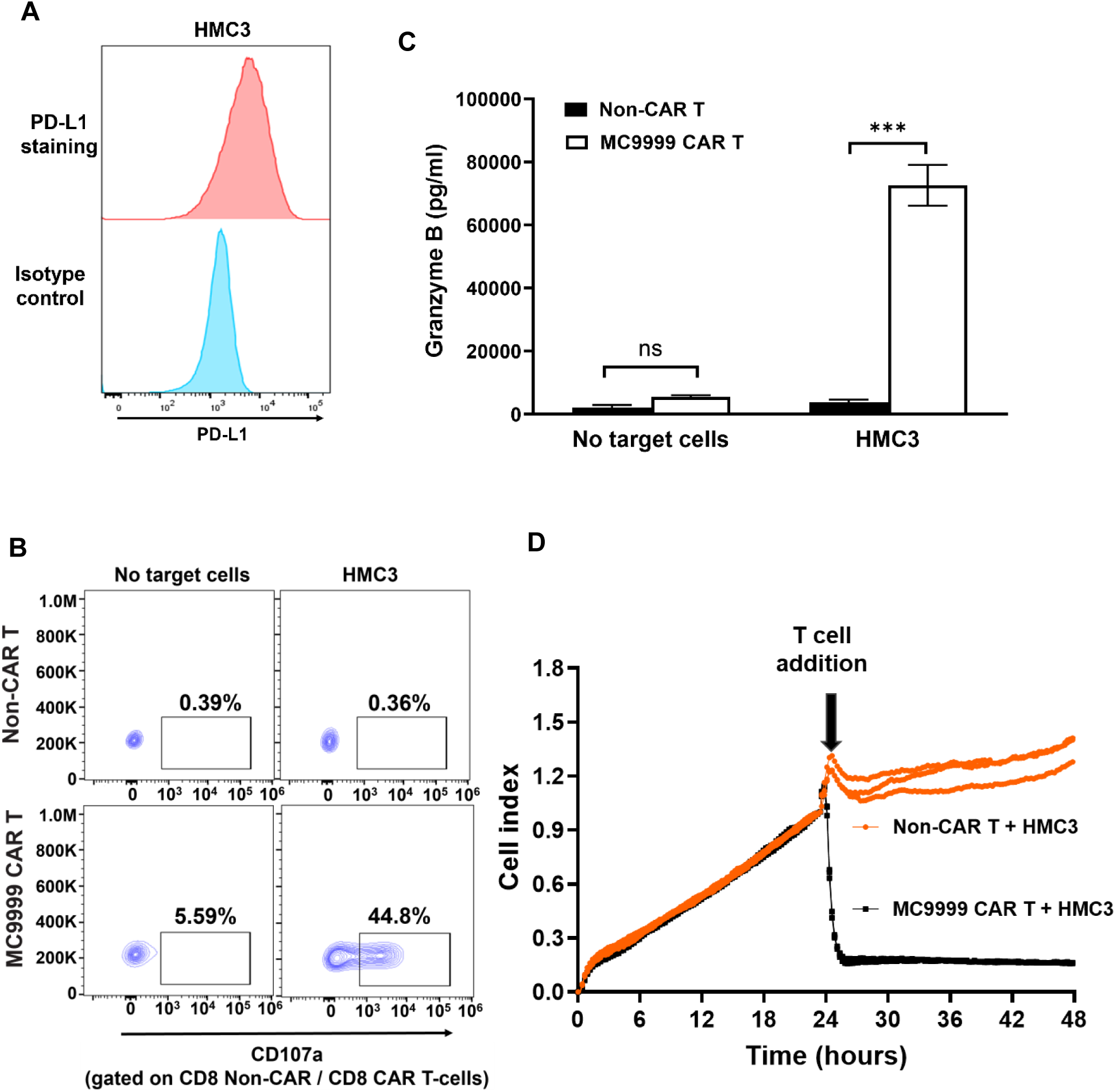
MC9999 CAR T-cells elicited cytolysis on HMC3 cells modeling tumor-associated microglias. (A) Immunostaining of HMC3 microglial cells with an anti-human PD-L1 monoclonal antibody demonstrated cell surface expression of PD-L1. An isotype control antibody served as a negative control. (B) Co-incubating MC9999 CAR T-cells with HMC3 cells triggered T-cell degranulation, as evidenced by the cell surface detection of CD107a. (C) A significant release of granzyme B was detected with ELISA when MC9999 CAR T-cells were co-incubated with HMC3 cells. The data, plotted as mean ± SEM of triplicate sampling, are representative of three independent experiments and analyzed using the multiple T test (*** p < 0.001). (D) The impedance-based killing assay revealed a real-time cytotoxicity exhibited by MC9999 CAR T-cells against HMC3 cells. HMC3 cells were cultured for 24 hours followed by addition of MC9999 CAR T-cells, which resulted in a significant decrease in the cell index, a measure of cellular impedance, within the cultured HMC3 cells. The data are representative of three independent experiments.

Immunosuppressive macrophages, including TAMs, typically exhibit an M2 phenotype within the TME. To model this population, we derived MDM-M2 macrophages and confirmed their MDM-M2 phenotype with the expression of CD163 and CD209 cell surface markers (Figure 7A, scatter plot). The expression of PD-L1 was detected on these M2 macrophages, distinguishing themselves from their CD14+ monocyte precursor (Figure 7A, histograms) and marking M2 macrophages as a target for MC9999 CAR T-cells (Figure 7B). This PD-L1 targeted-cytotoxicity was further confirmed with the significant release of granzyme B by MC9999 CAR T-cells upon interaction with M2 macrophages (Figure 7C). Moreover, the data from an impedance-based killing assay demonstrated the direct killing of M2 macrophages by MC9999 CAR T-cells (Figure 7D).

**Figure 7.**
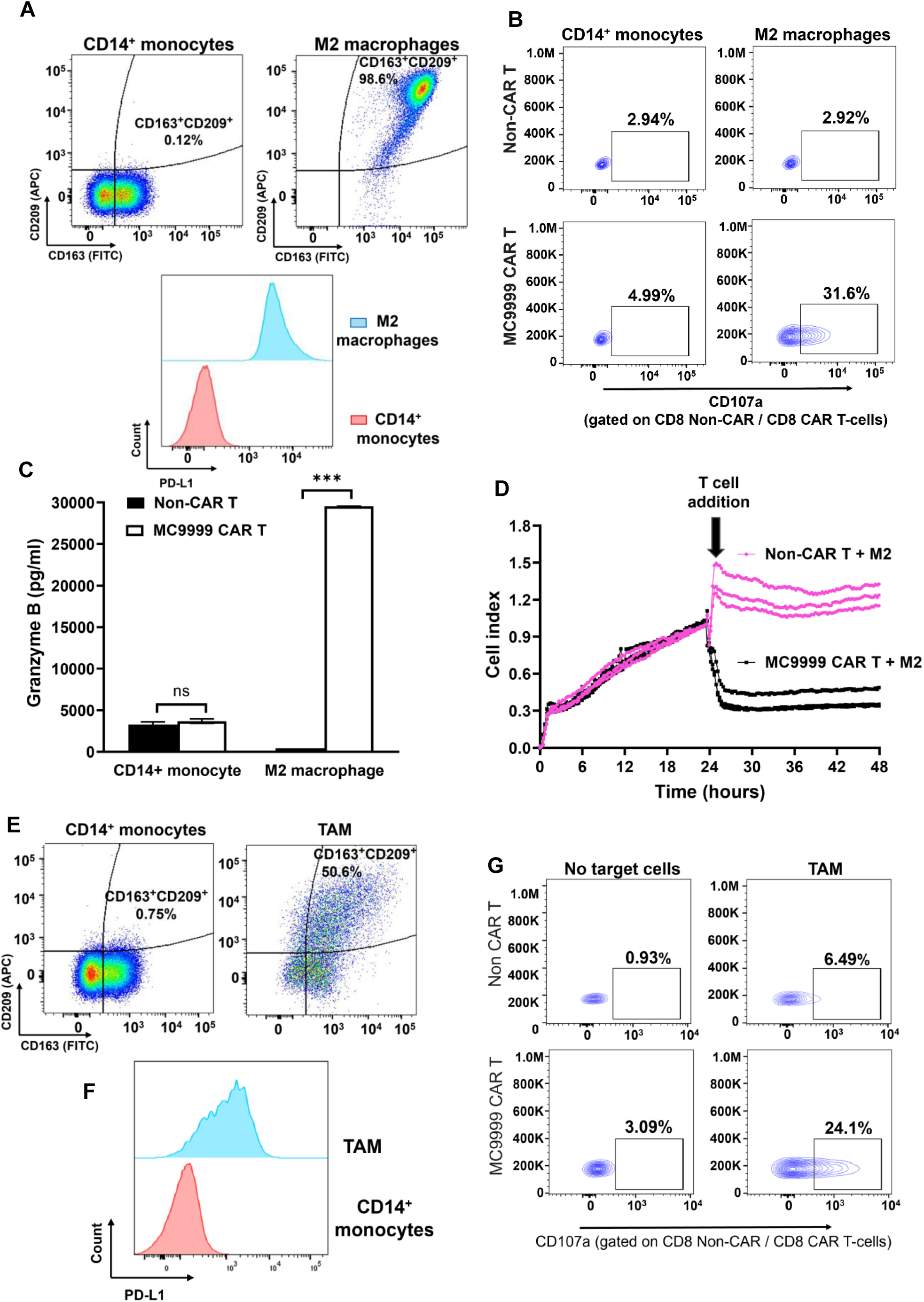
MC9999 CAR T-cells target monocyte-derived M2 macrophages that model immunosuppressive cells as well as patient-derived TAMs. (A) The CD163**^+^**CD209**^+^** immunophenotype of MDM-M2 macrophages was characterized and compared to that of their monocyte precursors (scatter plot). The PD-L1 expression on MDM-M2 macrophages was confirmed by immunostaining and analyzed using flow cytometry, with CD14**^+^**monocytes serving as a negative control (histogram plots). (B) MC9999 CAR T-cells exhibited cytotoxicity against MDM-M2 macrophages but not CD14**^+^** monocytes, as determined via a CD107a degranulation assay. (C) The release of granzyme B was significantly increased when MC9999 CAR T-cells were cultured with MDM-M2 macrophages. The data, plotted as mean ± SEM of triplicate sampling, are representative of three independent experiments and analyzed using the multiple T test (***, p < 0.001; ns, no significance). (D) The impedance-based killing assay demonstrated the direct killing of MDM-M2 macrophages by MC9999 CAR T-cells, as evidenced by a significant decrease in the cell index upon the addition of the CAR T-cells to cultured MDM-M2 macrophages. The data are representative of three independent experiments. (E) TAMs isolated from a surgically resected GBM tumor displayed the expected CD163**^+^**CD209**^+^**immunophenotype upon immunostaining. (F) The CD163**^+^**CD209**^+^**gated TAMs were highly positive for PD-L1 at the cell surface. CD14**^+^** monocytes served as the negative control for both immunophenotypic characterization and PD-L1 staining. (G) Evaluation via the CD107a degranulation assay revealed the MC9999 CAR T-cells, derived from healthy donor T cells, elicited cytotoxicity against the TAMs extracted from GBM tumor.

Finally, we assessed the cytotoxicity of MC9999 CAR T-cells on primary TAMs isolated from a surgically resected GBM tumor (QNS 960, Refer to Table S2). The immunophenotypic characterization verified the presence of a CD163+CD209+ double positive TAM population from the tumor (Figure 7E). Gating on this population, we identified PD-L1 expression on these immunosuppressive cells (Figure 7F). CAR T-cell degranulation was observed against TAMs (Figure 7G), underscoring the potential of MC9999 CAR T-cell therapy in targeting TAMs and subsequently mitigating immunosuppression within the TME of solid tumors.

### GBM patient-derived MC9999 CAR T-cells targeted primary GBM tumor cells

We have shown the antigen specific cytotoxicity of MC9999 CAR T-cells against a variety of PD-L1 expressing target cells that included both cancer cells and tumor-associated immunosuppressive cells. In these proof-of-principle studies, the CAR T-cells were generated from healthy donor T cells. Recognizing that T cell fitness in cancer patients may be compromised, we evaluated the cytotoxicity of patient-derived MC9999 CAR T-cells (from our GBM patients) against PD-L1-expressing target cells to highlight the translational significance. Following our laboratory SOPs for CAR T-cell production, we generated MC9999 CAR T-cells using peripheral blood T cells obtained from three GBM patients (Refer to Table S2). All three batches of patient-derived CAR T-cells exhibited acceptable quality control criteria with fold expansion (expansion fold > 25; Refer to Figure S3A) and the specific CAR T-cell characteristics of identity (identity >70%, CD3 staining; Refer to Figure S3B) and potency (potency >10%, tEGFR staining; Refer to Figure S3C).

The antigen specific cytotoxicity of these patient-derived MC9999 CAR T-cells was evaluated against the QNS120 and QNS712 patient-derived tumor cell lines, as well as the HMC3 cell line, with the corresponding Non-CAR T-cells as a negative control. The CAR T-cells derived from patient 1 and patient 2 exhibited comparable degranulation activity against the target cells, whereas those from patient 3 showed less activity (Figure 8A). In a separate measure of activity, we examined the release of cytotoxic granules and cytokines by the patient-derived MC9999 CAR T-cells in response to target cells and detected granzyme A, granzyme B, IFN-γ, and perforin in all cases (Figure 8B). As expected, variations were observed among the CAR T-cells derived from different patients. CAR T-cells from patients 1 and 2 released greater amounts of cytotoxic granules and IFN-γ than those from patient 3, aligning with the degranulation results. Our generation of three batches of qualified, functional patient-derived MC9999 CAR T-cells shows the translational feasibility of these CAR T-cells for clinical applicability.

**Figure 8.**
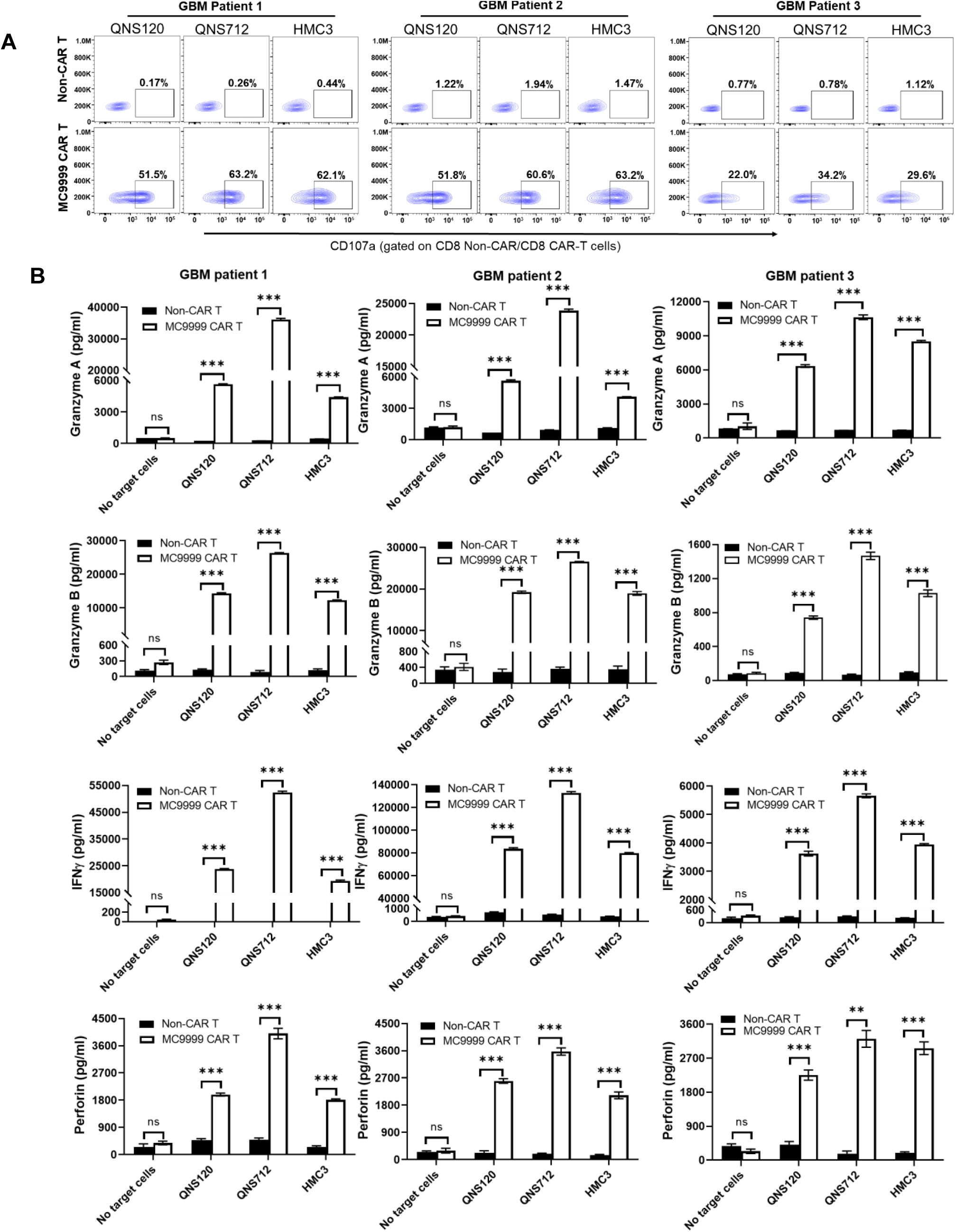
Validation of cytotoxic functionalities of GBM patient-derived MC9999 CAR T-cells. (A) Using peripheral blood T cells obtained from GBM patients, three batches of patient-derived MC9999 CAR T-cells were generated. The cytotoxicity of these patient-derived CAR T-cells was evaluated through a CD107a degranulation assay. Upon incubation with the PD-L1-expressing target cells including QNS120 and QNS712 GBM patient-derived tumor cells, as well as HMC3 microglia cells, the CAR T-cells exhibited degranulation activities, as evidenced by the presence of CD107a at the cell surface. (B) After 72 hours of co-culturing patient-derived MC9999 CAR T-cells with the target cells, the tissue culture supernatant was collected for quantitative analysis of T-cell cytotoxic granules/cytokines, including granzyme A, granzyme B, IFN-γ, and perforin, using a customized U-PLEX Multiplex Assay. The data were plotted as mean ± SEM of triplicate sampling and analyzed using the multiple T test (*** p < 0.001; ** p = 0.0011).

## DISCUSSION

To address challenges of CAR T-cell effectiveness in solid tumors, we have engineered PD-L1 targeting MC9999 CAR T-cells and have shown cytotoxicity and antitumor effects against various PD-L1 expressing tumor cells and macrophages that modulate the TME. The underlying rationale was that both tumor cells and immunosuppressive cells exploit PD-L1 pathways to evade immune surveillance. To this aim, we employed a humanized monoclonal antibody against human PD-L1^12^ to develop a PD-L1 CAR construct, MC9999. Using the MDA-MB-231 PD-L1 OE and MDA-MB-231 PD-L1 KO pair of cell lines, we validated antigen-specific cytotoxicity and antitumor effects of MC9999 CAR T-cells in both in intro and in vivo settings. Furthermore, MC9999 CAR T-cells exhibited activity against a diverse set of PD-L1 expressing, solid tumor-derived target cells that included a NSCLC cell line (Calu-1), a melanoma cell line (SH-4), a GBM cell line (LN229), and, finally, two patient-derived GBM cell lines. The observation that IV dosed MC9999 CAR T-cells eradicated intracranially established GBM tumors is highly encouraging for the future translational application of MC9999 CAR T-cell therapy for patients with GBM. Once high expression of PD-L1 was confirmed in three immunosuppressive macrophage models, we also showed the cytotoxicity of MC9999 CAR T-cells against HMC3 microglial, M2-macrophages, and patient-derived TAMs. These proof-of-principle studies highlight the potential effectiveness of targeting PD-L1 with MC9999 CAR T-cells against solid tumors.

Immunotherapies, specifically CAR T-cells and ICIs, mobilize the patient’s immune system to battle cancer. Indeed, CAR T-cell therapies have revolutionized the treatment of hematological cancers with complete remissions reported as high as 71–81% for relapse/refractory(r/r) acute lymphoblastic leukemia.^15^ This success story inspires researchers to continue the quest of a CAR T-cell therapy for treatment of solid tumors, even though this success has yet to be fully realized. Challenges remain, specifically with solid tumors having heterogeneous antigens or lacking a restrictive antigen, and there are challenges of trafficking into tumor and retaining activity of CAR T-cells once in the immunosuppressive TME.^16^ The identification of the immune checkpoints that regulate T cell function birthed ICIs, with a prominent targeted pair of PD-1 and its ligand, PD-L1.^1^ We were particularly drawn to the PD-1/PD-L1 cascade with the well characterized relationship between increased PD-1 expression and suppressed T cell function along with the observation that tumor cells and regulatory/support cells both can express PD-L1 to evade T-cell-mediated antitumor immunity. The presence of PD-L1 is a negative prognostic marker^17,18^ with reports of circulating PD-L1 positive monocytes, associated with some cancers.^19–21^ Hence, we hypothesized that designing PD-L1 targeting CAR T-cells would not only be able to strike tumor cells but would also attack the PD-L1 expressing regulatory/support cells resulting in the destabilization of the cellular network that generates the immunosuppressive TME.

This immunosuppressive TME has a cellular composition that forms a cellular network in which regulatory and support cells control access, modulate activation, and suppress activity of immune cells, specifically cytolytic T cells and tumor infiltrating lymphocytes (TILs) that are tasked with eradicating tumor cells.^22^ Initially classical immunohistochemistry and more recently advanced methods like spatial transcriptomics are contributing to the representative image of solid tumors^23–25^ with macrophages sometimes occupying more than 50%.^26^ Elevated PD-L1 levels have been consistently observed in various immunosuppressive cells within the TME, including TAMs, MDSCs, and even Tregs, which makes PD-L1 a viable therapeutic target for these immunosuppressive cells. With this perspective in mind, the MC9999 CAR was uniquely engineered to target not only tumor cells but also immunosuppressive cells within the TME for more effective therapeutic outcomes. We tested our hypothesis by validating the cytotoxicity of MC9999 CAR T-cells against three immunosuppressive macrophage models that highly express PD-L1, including HMC3 microglial cells, MDM-M2 macrophages, and patient-derived TAMs. These proof-of-principle studies highlighted the potential effectiveness of targeting PD-L1 with MC9999 CAR T-cells against the TME.

The success of PD-1/PD-L1 ICIs for the treatment of malignancies is remarkable, and the generation of a PD-L1 targeted CAR T-cells that allows for a more permanent elimination of PD-L1 expressing target cells remains a goal. Our findings solidify MC9999 CAR T-cells as a valid immunotherapy option with broad application to solid tumors with activity against not only tumor cells but also immunosuppressive cells within the TME. Generating MC9999 CAR T-cells from the peripheral T cells of GBM patients is highly encouraging particularly after measuring their activity against both microglial cells and patient derived GBM cell lines. We are actively developing a novel approach that leverages the operating room for intratumoral delivery of CAR T-cells, which allows effective T-cell application directly to tumor sites. We are encouraged with the response of MC9999 CAR T-cells against diverse cell lines since GBM, NSCLC, and melanoma have antigen heterogeneity while also expressing PD-L1 to induce immunosuppression.^27^

Despite these promising proof-of-concept data, we also acknowledge several remaining challenges before our goal can be actualized. Perhaps the most concerning issue related to any PD-L1 targeted CAR T-cell is the safety profile, particularly the risk of on-target, off-tumor effects that may lead to adverse clinical events. To mitigate these potential risks, we are optimizing the safety of MC9999 CAR T-cell therapy with strategies that include the localized as opposed to systemic CAR T-cell delivery to minimize potential side effects as well as upgrades to our CAR design. We are also exploring the feasibility of intracranial delivery of MC9999 CAR T-cells during neurosurgery for the treatment of GBM. Our current MC9999 CAR construct incorporates tEGFR as a safety switch, enabling the depletion of CAR T-cells upon introduction of cetuximab. Another strategy related to CAR construct design is our modeling and engineering of low affinity variants of our PD-L1 antibody that will spare normal tissues with low levels of antigen expression yet retaining binding affinity to target tumor cells that overexpress antigen. Lastly, we are also considering a SynNotch CAR design strategy^28,29^ to couple our MC9999 CAR with a more tumor-specific CAR, aiming to enhance tumor-specific targeting of MC9999 CAR T-cells. These safety-focused strategies will aid in the translation of a safe MC9999 CAR T-cell therapy into clinical applications for treating solid tumors.

## Materials and methods

### Cell lines and culturing conditions

The cell lines of MDA-MB-231, Calu-1, SH-4, LN229, HMC3, 293FT, and Jurkat were purchased from American Type Culture Collection (Manassas, Virginia, USA) and maintained in either 90% RPMI 1640, Iscove’s MDM, or 90% Dulbecco’s Modified Eagle Medium (Thermo Fisher, USA) supplemented with 10% heat-inactivated fetal bovine serum (Thermo Fisher, USA). Cell lines were authenticated by flow cytometry. The PD-L1 over-expression variant cell lines of MDA-MB-231 PD-L1 OE, LN229 PD-L1 OE, and SH-4 PD-L1 OE were generated as previously described^13^. PD-L1 knock out was induced in using IDT’s CRISPR-Cas12a (Cpf1) format involving ribonucleoprotein (RNP) complex between *Acidaminococcus* (As) Cas12a enzyme and PD-L1 gRNA (TATTCATGACCTACTGGCATT) targeting exon-2 of PD-L1 following IDT product instructions. For PD-L1 knock out in MDA-MB-231, LN229, Calu-1, and SH-4, the following nucleofector® 4D programs were chosen CH125, DS138, EO120, and EH100, respectively. Knockout cell lines were established from single cell clones post flow-sorting. Luciferase expressing human cell lines for in vivo experiments were generated as described.^13^ Prior to cryopreservation, the antigen-specific cell lines were authenticated using flow cytometry. All cell lines were routinely tested for mycoplasma contamination.

### PBMCs and Tn/mem isolation from healthy donors blood samples

Peripheral blood mononuclear cells (PBMCs) were obtained from healthy volunteer donors via leukapheresis using leukocyte reduction system (LRS) cones, by the Division of Transfusion Medicine, Mayo Clinic, Rochester, Minnesota, following current regulatory requirements and as previously described.^30^ To generate CAR T-cells, naïve and memory T cell (Tn/mem) populations were isolated from PBMCs in a three-step procedure, involving negative selection of both CD14 and CD25, followed by positive selection of CD62L, using CD14, CD25, and CD62L microbeads, adhering to the manufacturer’s protocol (Miltenyi Biotech, Germany).

### Isolation of T cells from blood samples of patients with GBM

The GBM patient blood procurement was performed under the biorepository protocol (IRB#17-003013) approved by the Mayo Clinic, Florida Institutional Review Board (IRB). All patients provided written informed consents, and the protocol adhered to the ethical principles of the Declaration of Helsinki. A single approximate 20 ml blood sample was collected. Disease-characteristics of these patients were recorded (Table S2). T cells were isolated using the Pan T cell isolation kit (Miltenyi Biotec, Germany).^13^

### CAR T-cell generation

A second-generation PD-L1-CAR (MC9999) was designed consisting of a novel PD-L1 antibody scFv,^12^ a hinge region with a CD4 transmembrane domain, as well as 4-1BB and CD3ζ intracellular signaling domains, complemented with a truncated EGFR (Figure S1). The CAR cDNA was integrated into pHIV.7 lentiviral vector. To ensure efficient lentivirus production, we utilized 293FT cells, followed by concentration and titer determination using Jurkat cells.

Tn/mem or pan-T cell populations, isolated from PBMCs, were divided into two aliquots for generating Non-CAR T-cells as control and another for generating CAR T-cells. The detailed CAR T-cell production protocol and accompanying quality control assays follow established methods.^13^

### Brain tumor initiating cell (BTIC) isolation and expansion from tumor tissue

Patient tumor tissue was collected with informed consent and approved by the Mayo Clinic, Florida Institutional Review Board (IRB# 16-008485). BTICs were isolated and expanded from primary patient glioblastoma tumor samples, and they were cultured under normoxic conditions (37°C, 5% CO_2_, 20% O_2_) following previously established methods.^31–37^ QNS120 and QNS712 are two such cell lines. In brief, the process involved the intraoperative resection of GBM tumor tissue, which was immediately delivered to the lab for processing. The tissue was mechanically dissociated followed with enzymatic digestion using TrypLE-express (Thermo Fisher, USA) to separate tumor cells from other tissue. The separated cells were then passed through a 40 μm nylon mesh, collected in a 50 ml conical tube, and centrifuged at 200xg for 5 minutes. After aspirating the supernatant, the cell pellet was resuspended in 1ml of stem cell media consisting of DMEM/F12 (Gibco, USA), 1% v/v Anti-anti (Sigma, USA), 2% v/v Gem21 NeuroPlex™ Serum-Free (without Vitamin A) (Gemini Bio-Products, USA), FGF (20 ng/ml PeproTech, USA), and EGF (20 ng/ml PeproTech, USA). After re-suspending, cell count and viability were measured using the Vi-Cell™ XR Cell Viability Analyzer (Beckman Coulter, USA). The cells were seeded as single cell suspension in non-adherent culture flasks at a density of 1.2 x10^4^ cells/cm^2^. These cells were cultivated as oncospheres for three passages before being plated and expanded on laminin-coated flasks for the establishment of cell lines. Throughout this process, all cells were maintained in stem cell promoting media containing EGF and FGF.

### Monocyte derived M2 macrophages from PBMCs

CD14-positive monocytes were isolated using positive selection with CD14 microbeads (Miltenyi Biotech, Germany). Macrophages were generated from this CD14-positive population by activation with M-CSF (25 ng/ml, PeproTech, USA) over a 7-day period. On day 7, M2 polarization was induced by the addition of Interleukin 4 (20 ng/ml, PeproTech, USA), in conjunction with M-CSF, for an additional 48-72 hours.^38^ Following M2 polarization, the cells were harvested, and their phenotype and PD-L1 expression were subsequently assessed via Fortessa flow cytometry (BD, USA) by staining for BUV395-CD14(clone M5E2, BD Horizon, USA), APC-CD209 (clone DCN46, BD Pharmingen, USA), AF488-CD163 (clone MAC2-158, BD Pharmingen, USA), and BV650-PDL-1 (clone29E.2A3, Biolegend, USA).

### Isolation of TAMs from tumor tissues of patients with GBM

GBM patient tumor tissue was collected with informed consent (IRB# 16-008485). The tumor tissue was first mechanically dissociated and further processed with the Tumor Dissociation Kit (cat # 130-095-929, Miltenyi, Germany) and the gentleMACS^™^ Dissociators (Miltenyi Biotech, Germany) for single cell dissociation. The separated cells were then passed through a 40μm nylon mesh, collected in a 50 ml conical tube, and centrifuged at 300xg for 7 minutes. After aspirating the supernatant, the cell pellet was resuspended in 10 ml PBS for lymphocyte cell isolation using a Ficoll gradient centrifuge, following previously established methods.^13^ T cells were removed by negative selection, and the remaining cells were analyzed via Fortessa flow cytometry (BD, USA). The analysis included staining for BUV395-CD14 (clone M5E2, BD Horizon, USA), APC-CD209 (clone DCN46, BD Pharmingen, USA), AF488-CD163 (clone MAC2-158, BD Pharmingen, USA), and BV650-PDL-1 (clone29E.2A3, Biolegend, USA).

### In vitro functional assays Degranulation assays

CAR T-cells were incubated with target cells at an effector-to-target (E:T) ratio of 2:1 in complete RPMI 1640 medium containing GolgiStop™ Protein Transport Inhibitor Reagent (BD Bioscience, USA) and CD107a APC antibody (BD Biosciences, USA) for 6 hours. The cells were subsequently stained with anti-CD3 BV605 (BD Biosciences, USA), anti-CD4 PE-Cy7 (BD Biosciences, USA), anti-CD8 APC-Cy7 (BD Biosciences, USA), and anti-EGFR BV421(BD Biosciences, USA). Samples were evaluated using the Attune flow cytometer (Thermo Fisher Scientific, USA) or the Fortessa flow cytometer (BD Biosciences, USA), and data were analyzed using FlowJo Version 10 software. Non-CAR T-cells from the same donors were used as negative controls.

### Granule release assay

CAR T-cells and target cells were co-incubated for 72 hours at an E:T ratio of 4:1. After the incubation period, the supernatant was collected and evaluated for granule protein release. The levels of granule proteins involved in cytotoxic T cell activity, specifically granzyme A, granzyme B, interferon (IFN) γ, and perforin, were quantified using a customized U-PLEX Multiplex Assay kit, following the manufacturer’s instructions (Meso Scale Diagnostics, Rockville, MD, USA.

### Impedance-based tumor cell killing assay (xCELLigence)

All experiments were performed using the respective target cell culturing media. The seeding of target cells was performed in 100 μl of medium per well to E-Plates 96 (Roche, Grenzach-Wyhlen, Germany), and appropriate cell densities were determined through titration experiments. Cell attachment was continuously monitored using the RTCA SP instrument (Roche) and RTCA software Version 1.1 (Roche) until the plateau phase was reached, typically occurring after approximately 24 hours. Subsequently, T cells were introduced at E:T ratio of 40:1 in 100 μl of medium. Impedance measurements were measured every 15 minutes for a duration of up to 96 hours. Each experiment was conducted in triplicate. Changes in electrical impedance were quantified as a dimensionless cell index (CI) value, which is derived from relative impedance changes corresponding to cellular coverage of the electrode sensors, normalized to baseline impedance values with medium only. For data analysis, CI values were exported, and the percentage of lysis was calculated relative to the control cells alone.

### Animal studies with bioluminescence imaging

NOD scid gamma (NSG) mouse breeding pairs were purchased from The Jackson Laboratory (stock no. 005557) to establish a breeding colony that was monitored in a pathogen-free animal facility at the Animal Resource Center at Mayo Clinic Florida, per institutional guidelines. Animal studies were approved by and in accordance with guidelines of the Institutional Animal Care and Use Committee (IACUC: 15020; protocol number A00005759 and A00006674). Mice (8-12 weeks old) received an intravenous (IV) ^39^ with a luciferase-expressing human tumor cell line (optimized in a separate experiment), randomized into test groups (5 mice per group), and treated with a single IV treatment dose of one of three treatments: PBS, Non-CAR T-cells, or CAR T-cells (concentrations in specific figure legends). For the GBM model, mice (8-12 weeks old) received an intracranial challenge^34,37^ with a luciferase-expressing human tumor cell line (optimized in a separate experiment), randomized into test groups (5 mice per group), and had two IV treatments spaced one week apart; treatment was either PBS, Non-CAR T-cells, or CAR T-cells. The tumor burden was quantified weekly by bioluminescent signal intensity on isoflurane-anesthetized mice that received a subcutaneous injection of D-luciferin (150 μg luciferin/1 g mouse body weight) 10 min prior to IVIS® imaging (PerkinElmer, Waltham, MA,USA). Survival data were presented and reported in Kaplan–Meier plots.^39^

## Acknowledgements.

We would like to acknowledge the funding support to HQ, which includes the Florida Health Grant (#MOG07, SB2500), the Mayo Clinic Florida CAR-T Manufacturing Program Fund, the Florida Department of Medicine Team Science Award, and the Mayo Clinic President’s Discovery Translation Program Award. The authors thank the Animal Resource Center at Mayo Clinic Florida for daily care of mice used in this study, the flow cytometry facilities at Mayo Clinic Florida and the Neurosurgery Biospecimens Repository of Intraoperative Patient Donations to Foster Collaborations across the Globe and Enterprise (BRIDGE-https://www.mayo.edu/research/labs/brain-tumor-stem-cell-research/research/neurosurgery-bridge-biobank). This publication was made possible through the support of the Distinguished Mayo Clinic Investigator Award (A.Q.-H.) and the William J. and Charles H. Mayo Professorship (A.Q.-H.), the Uihlein and Jacoby Neuro-oncology Research Fund (A.Q.-H.).

## Author contributions

HQ, YL, and AQ-H designed the project and studies. ESG, MMB, MJUN, AOL, and VKJ developed GBM patient-derived tumor cell lines and conducted in vivo studies on GBM models. YL, YQ, TH, and SL participated in the development of the MC9999 CAR construct, generating experimental models, performing CAR T-cell experiments, and analyzing resulting data. HQ, MEG, and YL created figures and prepared the manuscript. HD developed the anti-human PD-L1 mAb and contributed to the conception of the MC9999 CAR T-cell therapy. YL, TP, RD, MAK-D, and AQ-H contributed to disease-specific experimental design, data analysis, and manuscript review. Finally, HQ and AQ-H supervised the entire project.

## Competing interests

MAK-D declares research/grant from Bristol Myers Squibb, Novartis and Pharmacyclics. Consultancy for Kite Pharma. All other authors have no project-related conflict of interest to be claimed.

## Supplemental Material

### Supplemental Tables

**Table S1:**
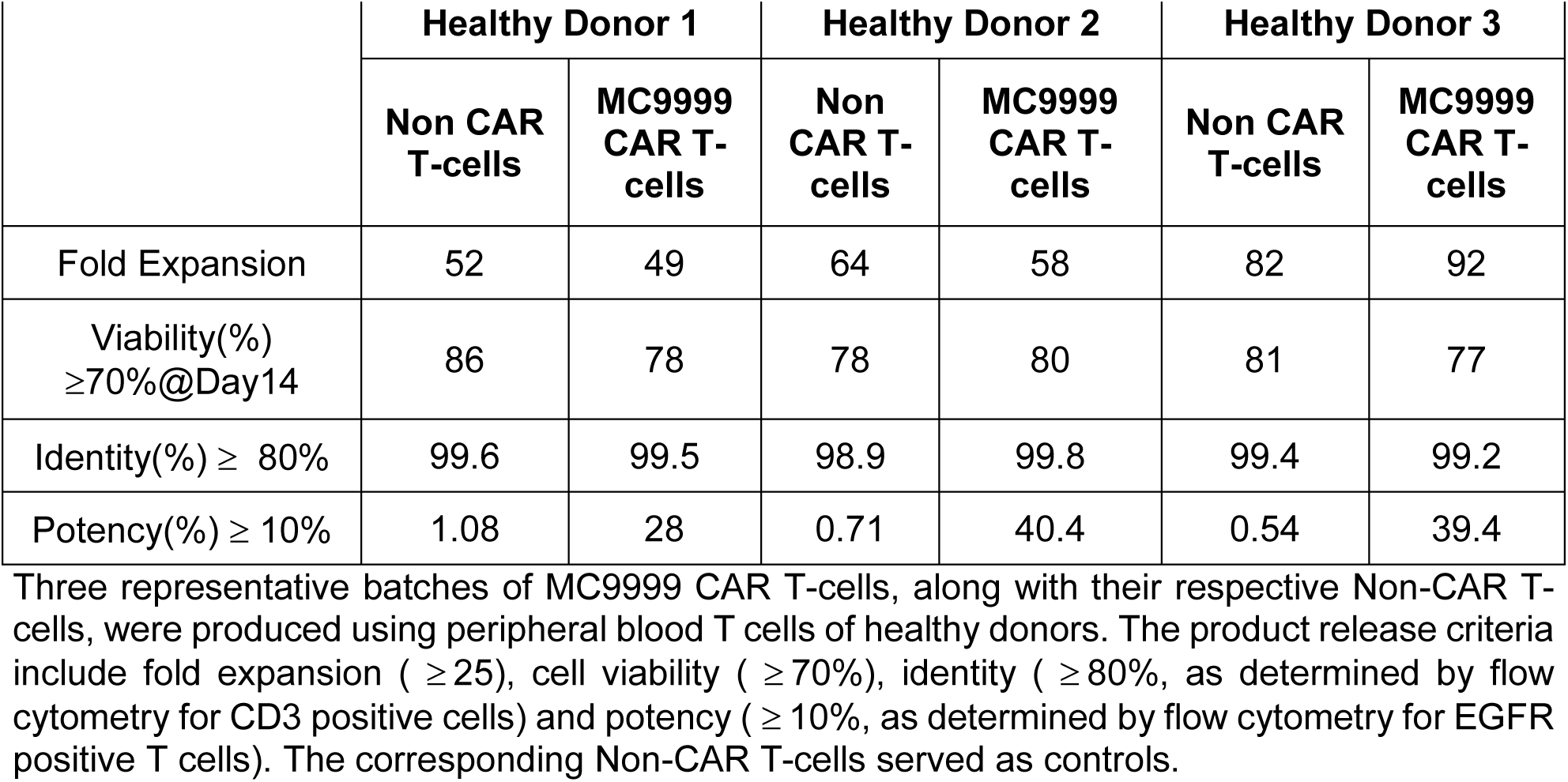
Characterization of representative batches of MC9999 CAR T-cells derived from healthy donors.

|  | Healthy Donor 1 |  | Healthy Donor 2 |  | Healthy Donor 3 |  |
| --- | --- | --- | --- | --- | --- | --- |
|  | Non CAR T-cells | MC9999 CAR T-cells | Non CAR T-cells | MC9999 CAR T-cells | Non CAR T-cells | MC9999 CAR T-cells |
| Fold Expansion | 52 | 49 | 64 | 58 | 82 | 92 |
| Viability(%)<br>≥70%@Day14 | 86 | 78 | 78 | 80 | 81 | 77 |
| Identity(%) ≥ 80% | 99.6 | 99.5 | 98.9 | 99.8 | 99.4 | 99.2 |
| Potency(%) ≥ 10% | 1.08 | 28 | 0.71 | 40.4 | 0.54 | 39.4 |

Three representative batches of MC9999 CAR T-cells, along with their respective Non-CAR T-cells, were produced using peripheral blood T cells of healthy donors. The product release criteria include fold expansion ( ≥ 25), cell viability ( ≥ 70%), identity ( ≥ 80%, as determined by flow cytometry for CD3 positive cells) and potency ( ≥ 10%, as determined by flow cytometry for EGFR positive T cells). The corresponding Non-CAR T-cells served as controls.

**Table S2:** General clinical information about the GBM patients.

| Laboratory-generated Patient ID | Patient age range | Diagnosis, disease stage | IDH-1/2 | MDMT methylation | Prior steroid use | Date of surgery | Past medical history | Onset symptoms | Figures |
| --- | --- | --- | --- | --- | --- | --- | --- | --- | --- |
| QNS120 | Patient in 50s | Primary Glioblastoma, Grade 4 | WT | Yes | No | 11/09/2017 | Atrial fibrillation status post ablation | Memory loss, fatigue, insomnia | Fig 4 A,B,C,D |
| QNS712 | Patient in 70s | Primary Glioblastoma, Grade 4 | WT | No | Yes (Dex) | 04/27/2021 | Atrial fibrillation status post ablation | Speech deficit, imbalance, confusion | Fig 4 A,B,C,D |
| QNS960 | Patient in 70s | Primary Glioblastoma, Grade 4 | WT | No | Yes (Dex) | 03/02/2023 | Scleroderma, Sjögren, primary biliary cholangitis | Seizures | Fig 7 A, B,C |
| GBM Pt 1 | Patient in 50s | Primary Glioblastoma, Grade 4 | WT | Yes | Yes (Dex) | 06/28/2023 | Diabetes, hypertension, asthma, obstructive sleep apnea, iron deficiency anemia | Headache | Fig 8 A,B,C and Supp Fig S4 |
| GBM Pt 2 | Patient in 60s | Primary Glioblastoma, Grade 4 | WT | Yes | Yes (Dex) | 06/28/2023 | Reflux, hyperlipemia, hypertension, gout, kidney stones | Memory problems, word searching issues, right arm numbness and tingling, and right-sided clumsiness | Fig 8 A,B,C and Supp Fig S4 |
| GBM Pt 3 | Patient in 60s | Primary Glioblastoma, Grade 4 | WT | Yes | No | 07/13/2023<br>07/20/2023* | Hypertension, diabetes, seizures, cataracts, rheumatoid arthritis | Disorientation, occipital headaches, reading and concentration issues | Fig 8 A,B,C and Supp Fig S4 |

Dex = Dexamethasone; \*Underwent repeat surgical resection one week after first resection.
Selected clinical information for the GBM patients who provided tumor tissue. Additional information tracks the patient samples that were used in the specific experiments and the corresponding figures.

### Supplemental Figures

**Figure S1.**
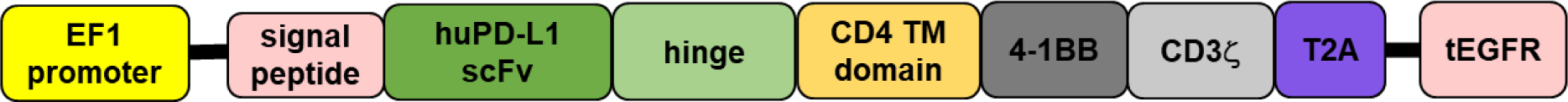
Schematic diagram of MC9999 CAR. The MC9999 CAR is composed of following elements in tandem: EF1 promoter, signal peptide, PD-L1 recognizing element (huPD-L1 scFv), hinge region, transmembrane domain (CD4 TM domain), costimulatory domain (4-1BB), intracellular T cell activation domain (CD3**ζ**), self-cleaving 2A peptide (T2A), and tEGFR (truncated EGFR that serves as a marker of CAR expression and a suicide switch mediated by cetuximab.

**Figure S2.**
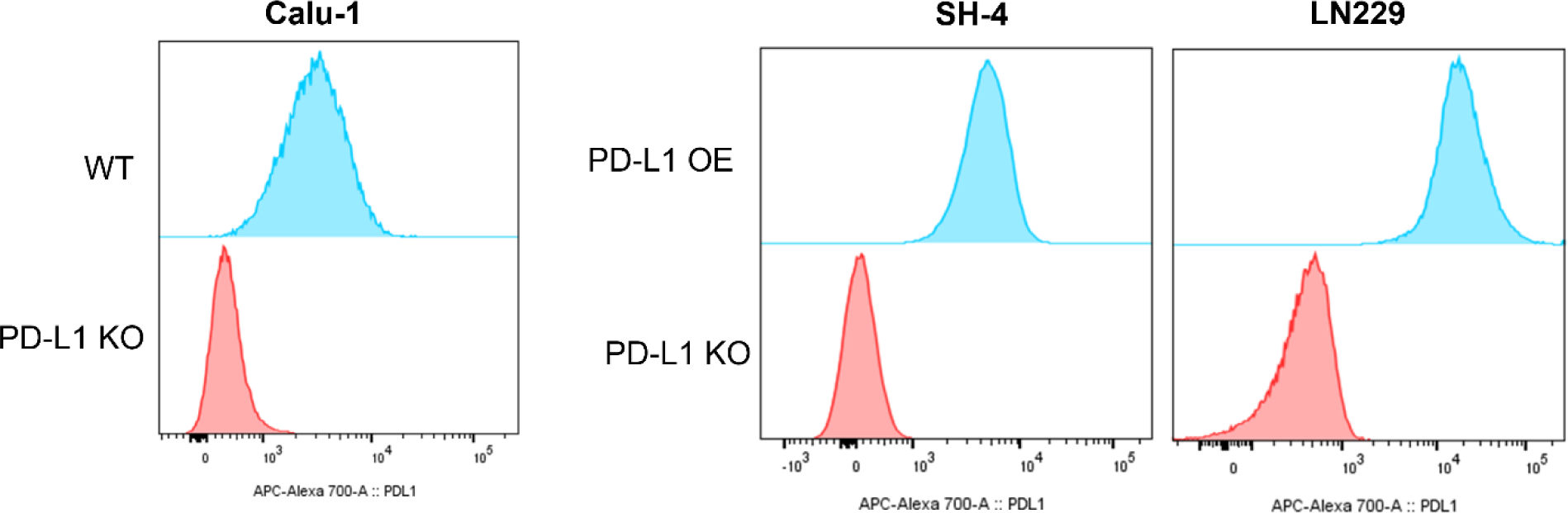
Characterization of PD-L1 expression in various solid tumor cell models. Calu-1 (NSCLC), SH4 (melanoma) and LN229 (GBM) cell lines were genetically modified to either lose (PD-L1 KO) or overexpress PD-L1 (PD-L1 OE). Validation of PD-L1 expression on these cell lines was conducted using a flow-based immunostaining assay.

**Figure S3.**
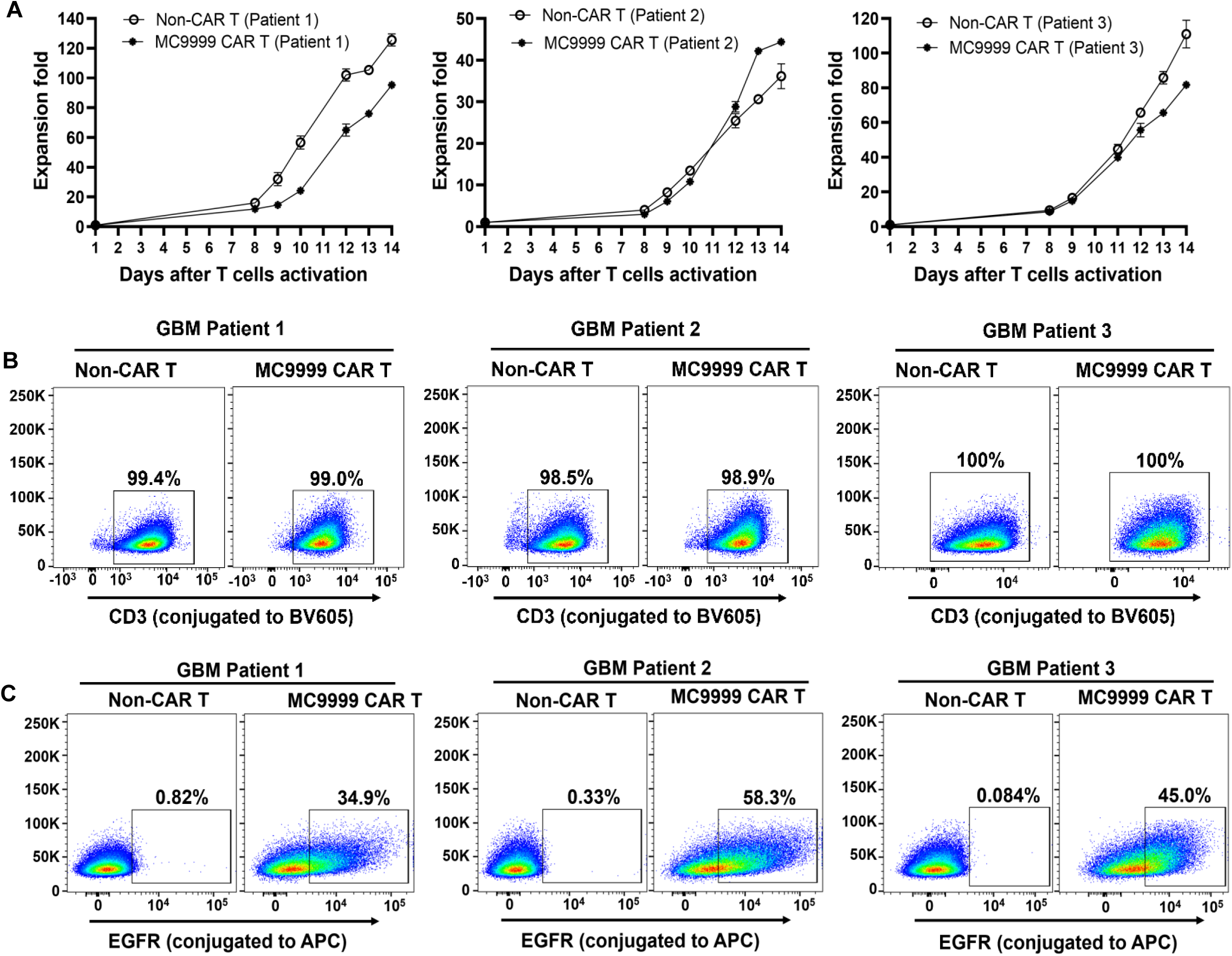
Characterization of the MC9999 CAR T-cells generated from three GBM patients. MC9999 CAR T-cells and their corresponding Non-CAR T-cells were derived from T-cells isolated from peripheral blood of three GBM patients. (A) The growth curves of CAR T-cells over 2-week expansion. Duplicated cell counts were collected for each time point. The produced CAR T-cells were immunostained for CD3 to confirm the identity (B), and EGFR to identify the potency (C).

